# Temporal-spatial transcriptomics reveals key gene regulation for grain yield and quality in wheat

**DOI:** 10.1101/2024.06.02.596756

**Authors:** Xiaohui Li, Yiman Wan, Dongzhi Wang, Xingguo Li, Jiajie Wu, Kunming Chen, Xue Han, Yuan Chen

**Affiliations:** National Key Laboratory of Wheat Improvement, Peking University Institute of Advanced Agricultural Sciences, Shandong Laboratory of Advanced Agriculture Sciences in Weifang, Weifang, Shandong, 261325, China; Key Laboratory of Plant Cell and Chromosome Engineering, Institute of Genetics and Developmental Biology, Chinese Academy of Sciences, Beijing 100101, China; National Key Laboratory of Wheat Improvement, College of Life Sciences, Shandong Agricultural University, Taian, Shandong, 271018, China; National Key Laboratory of Wheat Improvement, College of Agronomy, Shandong Agricultural University, Taian, Shandong, 271018, China; National Key Laboratory of Crop Improvement for Stress Tolerance and Production, College of Life Sciences, Northwest A&F University, Yangling, Shanxi, 712100, China; College of Plant Protection, Shandong Agricultural University, Taian, 271018, Shandong, China

**Keywords:** wheat, temporal-spatial transcriptomics, grain development, haplotype

## Abstract

Cereal grain size and quality are important agronomic traits in crop production. The development of wheat grains is underpinned by complex regulatory networks. The precise spatial and temporal coordination of diverse cell types is essential for the formation of functional compartments. To provide comprehensive spatiotemporal information about biological processes in developing wheat grain, we performed a spatial transcriptomics study during the early grain development stage from 4 to 12 days after pollination. We defined a set of tissue-specific marker genes and discovered that certain genes or gene families exhibit specific spatial expression patterns over time. Weighted gene co-expression network and motif enrichment analyses identified specific groups of genes potentially regulating wheat grain development. The embryo and surrounding endosperm specifically expressed transcription factor *TaABI3-3B* negatively regulates embryo and grain size. In Chinese breeding programs, a haplotype associated with higher grain weight was identified, linked to altered expression levels of *TaABI3-3B*. Data and knowledge obtained from the proposed study will provide pivotal insights into yield improvement and serve as important genetic information for future wheat breeding.

## Introduction

Wheat (*Triticum aestivum* L.) is the top three cereal crops worldwide, cultivated across approximately 230 million hectares globally. Enhancing wheat yield carries substantial implications for global food and nutrition security. The wheat grain comprises three primary components: the diploid embryo, the triploid endosperm, and the pericarp, also known as the seed coat (Chaudhury et al., 2001). Gene expression regulation in the seed coat, endosperm, and embryo undergoes spatiotemporal changes, forming functional compartments with distinct spatiotemporal characteristics. These tissues collaboratively regulate nutrient transport and grain development in wheat. However, current cell classifications of wheat grain are predominantly based on morphological observations and a limited number of specifically expressed genes (Zhong et al., 2023). Genetics study has helped to identify crucial gene involved in grain development (Rathan et al., 2022; Tekeu et al., 2021). Multi-omics analysis techniques are utilized to comprehensively understand the different composition of wheat grain (Khakimov et al., 2014; Zhang et al., 2021). All these studies typically lack the gene spatial information, limiting the understanding of the specific functions of each cell population.

Spatial transcriptomics offers single-cell level transcriptional information, enabling detailed exploration of tissue-specific gene expression (Bressan et al., 2023; Li et al., 2024a; Rhaman et al., 2024). Spatial transcriptomics has been extensively used in plant developmental stage study because it allows for the precise mapping of gene expression within the spatial context of developing tissues. In recent years, significant discoveries have been made in organogenesis, cell type identification, and cell fate lineage reconstruction through spatial transcriptomics (Giolai et al., 2019). In maize, spatial transcriptomics data identified genes specifically expressed in the apex of determinate meristems that are involved in stem cell determinacy (Wang et al., 2024). In peanuts, spatial transcriptomics indicated that stem-forming layer cells have the potential to differentiate into both xylem and phloem (Liu et al., 2022b). In orchids, the crucial roles of MADS-box genes and other potential downstream genes in organ initiation and morphogenesis have been elucidated through spatial transcriptomics (Liu et al., 2022a). By combine the temporal and spatial expression distribution of genes during maize kernels development constructed a comprehensive gene network regulating in grain filling process (Fu et al., 2023), By adding different germination time to barley germination spatial transcriptomics also help to reveal aquaporin gene families and auxin transport genes roles in zygote embryo activation and embryo polarity establishment processes (Peirats-Llobet et al., 2023). Meanwhile, integrating single-nucleus and spatial transcriptomics could enhance resolution of plant-microbe interactions in nodulating plants (Liu et al., 2023b). It can also localize rare or transitional cells within callus tissue, shedding light on their roles in processes in differentiation (Song et al., 2023). This integration allows for the creation of comprehensive gene expression maps and identify where specific genes are active and how their expression patterns change over time (Li et al., 2024b).

Grain size in wheat depends on signals from both the mother plant and the embryo itself, which work together to grow the embryo, endosperm, and seed coat. Similarly, in rice, different pathways control seed size, such as the ubiquitin-proteasome, G protein, and MAPK pathways, along with plant hormones and transcriptional regulators (Li et al., 2019). Transcription factors play crucial roles in the development of the embryo, endosperm, and seed coat. Some transcription factors specifically expressed in the endosperm and seed coat have been shown to regulate wheat grain development. For instance, the basic helix-loop-helix (bHLH) family transcription factor TaPGS1 is highly expressed in the aleurone layer cells and pericarp during early embryo development. By binding to the promoter of the transcription factor TaFl3, TaPGS1 influences wheat endosperm structure, regulating grain width, length, and thousand-grain weight (Guo et al., 2022). Another endosperm-specific transcription factor TaNAC019, interacts with the glutenin regulatory factor TaGAMyb, suppressing the expression of key starch synthesis genes *TaAGPS1-A1* and *TaAGPS1-B1*, thereby negatively regulating starch synthesis in the endosperm (Liu et al., 2020). TaABI19 binds to the promoter of the subgenome homologous gene of *TaPBF* and enhances its expression, affecting starch synthesis and grain development (Liu et al., 2023a).

The embryogenesis in wheat is controlled by a series of precise sequential transcriptional programs (Armenta-Medina et al., 2021; Dresselhaus and Jürgens, 2021). A crucial question is how signals from the embryo are swiftly transmitted to the endosperm, precisely regulating grain development by coordinating with endosperm-related genes. In maize, the transcription factor VIVIPAROUS-1 (VP1) is expressed in the scutellum, which links the endosperm and embryo. VP1 regulates genes related to globulin and nutrient metabolism pathways, facilitating the transfer of proteins from the endosperm to the embryo (Zheng et al., 2019). The embryo surrounding region (ESR) in wheat also plays a crucial role in signal transmission between the endosperm and embryo (Zheng et al., 2014). However, studies on the coordinated regulation of grain development by the embryo and endosperm in wheat are limited (Song et al., 2021). Understanding the key genes that regulate early embryonic development in wheat embryos and endosperm is essential.

Here, we use temporal-spatial transcriptomic analysis to create a detailed molecular spatial map, revealing specific spatial expression patterns during early grain development. From the spatial transcriptomic data, we identified a series of candidate genes and mutants involved in regulating wheat grain development. The functions of these key genes were further validated using genetic variations and transgenic materials. This study offers a valuable resource for understanding wheat grain development and provides new insights for enhancing yield.

## Results

### Generating spatial transcriptomes profiles of early development stage in wheat grain

To gain a comprehensive understanding of early grain development in wheat, we conducted spatial transcriptomics in Jimai 22, a variety widely cultivated in China for its high yield (Figure 1A). Three developmental stages were selected for cryosection: pro-embryo (4 days after pollination (dap)), transition (8 dap), and differentiation (12 dap) (Figure 1B). On day 4, early embryogenesis begins, forming the totipotent zygote, while the endosperm starts cellularization, and the pericarp begins accumulating starch granules. By day 8, the embryos and endosperm undergo rapid differentiation and expansion. By day 12, nutrient decomposition starts, with nutrients being transported to the endosperm and embryos. Longitudinal sections were performed at 4 dap, 8 dap, and 12 dap, with cross sections conducted specifically at 12 dap to better observe tissues such as cavity fluid.

**Figure 1.**
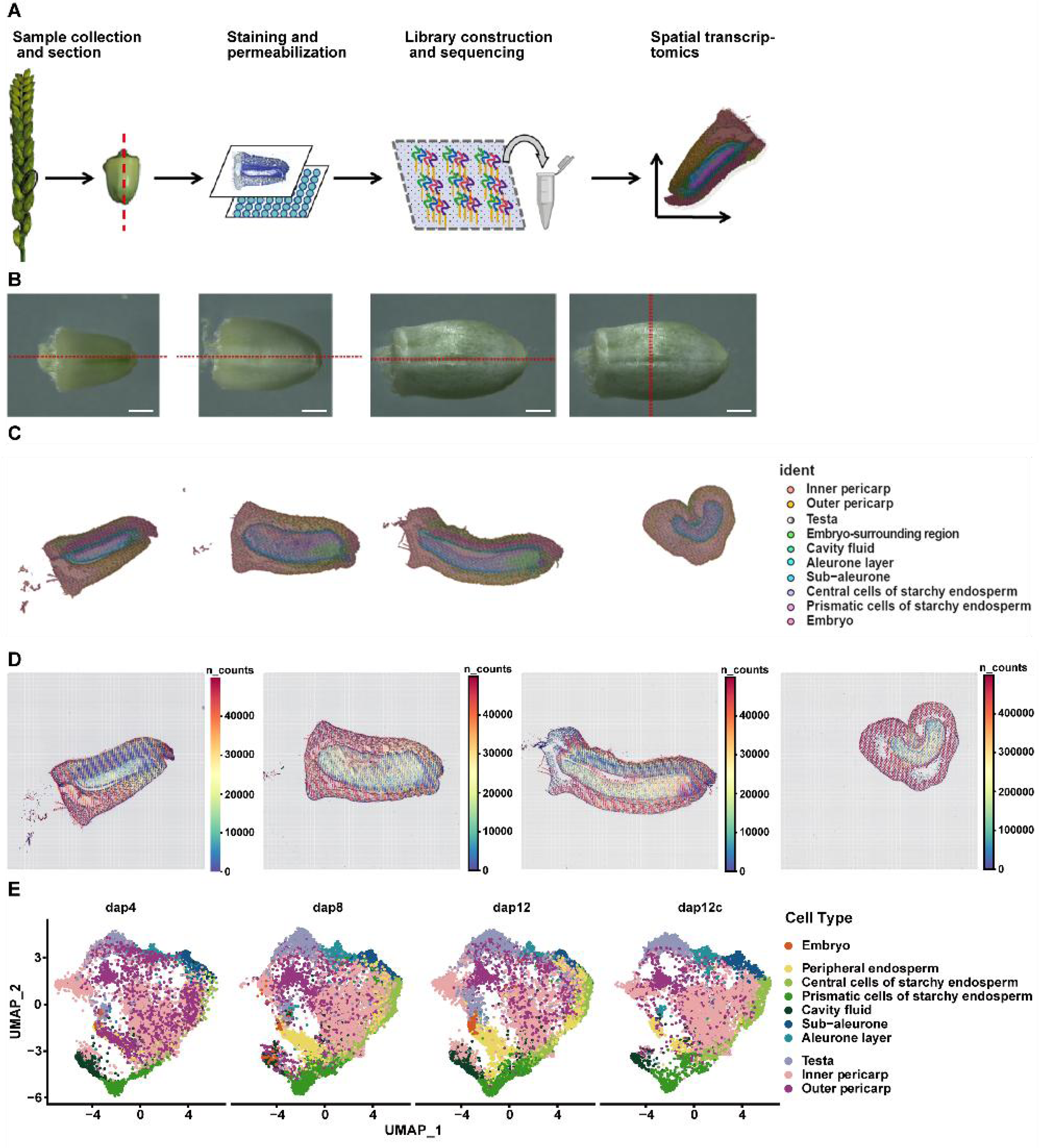
Spatially resolved transcriptome analysis of wheat grain. A. A workflow for sampling and sequencing of wheat grain on a BMKMANU S1000. B. Developing grain at 4, 8 and 12 dap. C. Spatial visualization of the unbiased spot clustering for 4, 8 and 12 dap wheat sections. Merged bright field image and spatial clusters of other three sections. The tissue/cell-type identity of each cluster was assigned based on the location of each cluster. D. Heat map and spatial distribution map of counts percent.mt expression in spot of each sample. E. Uniform manifold approximation and projection (UMAP) of spatial spots from 4, 8 and 12 dap wheat sections. Dots correspond to individual spots on the visium slide; colors indicate cluster association for each spot.

We prepared 10 μm thick sections and mounted them onto the sequencing areas of the BMKMANU S1000 Gene Expression chip (6.5 mm × 6.5 mm), which could fully cover the longitudinal and cross sections of the whole grain at the observed stages, facilitating the observation of gene expression across the entire grain. The chip contains 5000 spots arrayed with oligonucleotide barcodes to capture mRNA molecules, allowing transcripts to be mapped back to their original locations on the tissue to pinpoint their spatial distribution. Moreover, each slide contains approximately 2 million spots, serving not only for mRNA capture but also for determining spatial positional information. With a spot diameter of 2.5 μm and a spacing of approximately 5 μm between adjacent spots on the BMKMANU S1000 platform, this technology enables spatial transcriptomic analysis at subcellular resolution (Figure S1).

The cDNA library was sequenced using the BMKMANU S1000 platform (Figure S2). Unique Molecular Identifiers (UMIs) were utilized to quantify RNA molecules across the sample sections. Our results show that the number of tissue-covered spots across all sections ranged from 9,934 to 15,107, with a resolution of 27 μm, and the median UMI counts per spot varied from 330 to 1,786. Notably, a total of 15,107 spots were identified, with an average of 1,410 genes per spot (refer to Table S1 and Figure S3 A-B). Moreover, the genes identified in each section displayed a robust correlation (Spearman: r = 0.98-0.99), and sections of the same tissue type consistently clustered together (see Figure S3 C), indicating a high degree of data consistency.

We identified 12 spatial clusters based on gene expression similarities using dimensionality reduction. These clusters were present across all stages except for cluster 11 (Figure S4). By integrating anatomical information from cryosections and topuidine blue (TBO)-stained images, we delineated 10 functional cellular groups (Figure 1C), comprising: (1) two maternal regions - inner pericarp and outer pericarp; (2) seed coat region - testa; (3) embryo region; and (4) six endosperm regions: cavity fluid, aleurone layer, sub-aleurone, central cells of starchy endosperm, prismatic cells of starchy endosperm, and embryo-surrounding region (ERS).

We precisely delineated three interfaces: the cavity fluid, situated between maternal and filial tissues; the endosperm-ERS interface, located between the embryo and endosperm; and the endosperm-aleurone interface, existing between maternal and endosperm tissues. Spatial transcriptomics facilitates the simultaneous detection of RNA in situ expression data for approximately 80,000 genes across the entire wheat genome. This high-throughput capability enhances sensitivity and enables the discovery of novel cell types. Specifically, we identified six distinct cell types within the aleurone layer, five additional groups within the subaleurone layer, and nine groups within the cavity fluid (Figure S5).

The gene expression abundance across different tissues was statistically analyzed using the BSTMatrix software. The number of counts (nCounts) in spots was visualized through a heatmap (Figure 1D), while the number of unique molecular identifiers (nUMI) is presented in Figure 1E. According to the heatmap (Figure 1D), gene activity decreases in the outer layers, such as the pericarp and testa, as grain development progresses, but increases in the embryo and ESR. This suggests that the embryo gradually activates gene expression in nearby tissues. In the endosperm, the aleurone layer shows higher activity than the central endosperm, indicating that the aleurone layer is the first formed layer and begins to differentiate both inwardly and outwardly. This differentiation leads to the establishment of a specialized cell layer covering the entire surface of the endosperm, except in regions with transfer cells. Through UMAP clustering analysis of different chips, it was observed that various cell clusters exhibit consistent clustering across different developmental stages (Figure 1E). As the grain develops, the degradation of the pericarp and testa begins, and the endosperm gradually starts to proliferate and differentiate. At 8 dap, the number of endosperm cells significantly increases, with an even more pronounced increase at 12 dap, as indicated by the rising proportion of endosperm cells in the UMAP analysis.

In summary, our spatial transcriptomic sequencing of wheat grains partitioned the grains into 10 distinct cell clusters, revealing differential expression patterns and developmental modes across these clusters, provides a more detailed insight into the developmental processes of grains.

### Spatiotemporal analysis of key gene expression during wheat grains development

To verify that spatial transcriptomics can detect gene expression more accurately than other approaches, we compared our spatial transcriptomic data to a published embryo transcriptome analysis at 4 dap and 8 dap, as well as pericarp transcriptome data at the early leaf stage (Table S2). Notably, the embryo data could be confirmed by previous results, while endosperm and pericarp can be divided into new cell types based on spatial transcriptome data. According to laser microdissection RNA-seq data, a portion of marker genes from the embryo, endosperm, and pericarp are expressed in our data. We identified 12 spatial clusters through dimensionality reduction and clustering of spatial transcriptomic data using UMAP (see Figure 2A). Subsequently, clustering expression analysis was conducted between these 12 spatial clusters and the previously defined 10 cell clusters, revealing a correspondence between them (see Figure 2B). Powerful electronic RNA in situ hybridization was created based on the spatial transcriptomic data (Figure S6).

**Figure 2.**
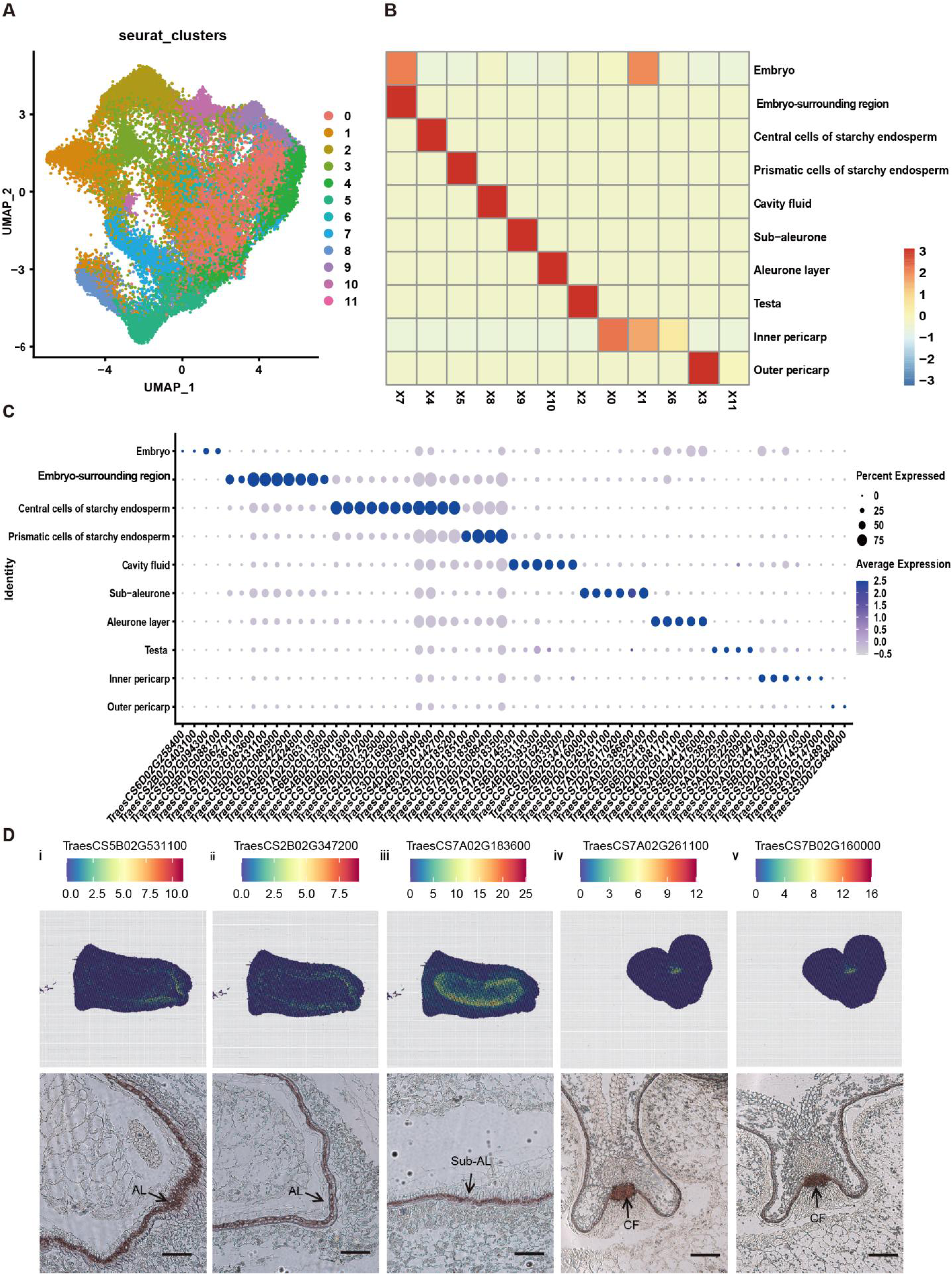
Identification of tissue-specific marker genes and their validation by in situ hybridisation. Identification and validation of tissue-specific marker genes using in situ hybridization. A. UMAP representation of the 12 clusters determined at the three timepoint. B. Heat map analysis of 12 clusters and 10 cell-types. C. Bubble plot showing transcript enrichment (average expression and percentage) of representative cell type-specific marker genes in the 10 cell-types. Spot colors correspond to the same tissues in the bubble plot. Genes highlighted in red were validated by in situ hybridization. D. In situ hybridization of selected marker genes (i) TraesCS5B02G531100; (ii) TraesCS2B02G347200; (iii) TraesCS7A02G183600; (iv) TraesCS7A02G261100 ; (v) TraesCS7B02G160000 confirmed localization of tissue-type specific transcripts at 12 dap timepoint. For each in situ hybridization, top panels show electronical RNA in situ hybridization and bottom panels with the antisense probes. Scale bar =100 μm.

Our spatial transcriptome data show that the endosperm can be divided into six cell groups. The aleurone layer, part of the endosperm, is rich in lipids and proteins, with its marker gene being the non-specific lipid-transfer protein TraesCS5B02G531100. The central cells of starchy endosperm, which forms last, primarily stores starch and sugars, with its marker gene being alpha-amylase TraesCS4B02G328100. Prismatic cells of the starchy endosperm store starch and sugars and are marked by seed storage-related proteins such as TraesCS1A02G063100. The cavity fluid includes transfer cells, essential for nutrient transport and storing some metal ions, with the marker gene being the sugar transporter SWEET, such as TraesCS7B02G160000.The pericarp can be divided into two cell types: outer and inner pericarp, which serve as nutrient storage tissue early in development and export nutrients during the filling stage. The lipid transfer protein TraesCS3A02G344700 marks the pericarp. The immature caryopsis exhibits a photosynthetically active testa, rich in chloroplasts and genes associated with photosynthesis, with TraesCS5B02G353200 as the marker gene (Figure S7). For each cell type, specific marker genes were identified based on tissue-specific expression and functional relevance (Table S3). In summary, seeds exhibit distinct clustering characteristics that closely align with their anatomical structure.

To validate the mRNA distribution, we conducted in situ hybridization experiments on five genes in different cell types. Consistently with spatial transcriptomic data (Figure 2D), the marker genes demonstrated specific expression localized to the aleurone layer, subaleurone layer, and endosperm cavity.

### Transcriptomic difference between grain cell type

To gain deeper insight into the spatial and temporal dynamics of grain filling and grain development, we divided the grain into five key components: seed coat, endosperm, embryo, aleurone layer, and cavity fluid (Figure 3A) for thorough analysis. We created co-expression networks for 10 seed cell types across three developmental stages, and then calculated the average expression of each gene within each cell type at every time point. A total of 11,365 eligible genes were grouped into 21 modules (Figure 3B, Table S4). The largest module, ME1, containing over 55.7% (6,332) of the genes, is mainly expressed in transport tissues like the aleurone layer, subaleurone layer, and endosperm cavity. Module ME2 comprises 1,181 genes primarily expressed in embryos at 4 dap, forming an embryo-specific expression group with ME19. Genes in the ME2/19 group are enriched in nucleocytoplasmic transport and amino acid biosynthesis pathways, as revealed by Kyoto Encyclopedia of Genes and Genomes (KEGG) pathway analysis (Figure 3C). Modules ME3/7/9/13/14/16 are associated with starch synthesis and nutrient accumulation in the endosperm, mainly enriched in starch and sucrose metabolism pathways. The aleurone layer is mainly composed of modules ME1/11/12/15, with genes primarily enriched in ribosome biogenesis, spliceosome assembly and protein processing in the endoplasmic reticulum. Conversely, the endosperm cavity consists of modules ME5/8/18, showing enrichment in pathways such as endocytosis, biosynthesis of secondary metabolites, and various metabolic pathways. Similarly, the seed coat primarily comprises modules ME0/6/17, enriched in pathways such as carbon fixation in photosynthetic organisms and pathways associated with photosynthesis and photosynthesis-related antenna proteins (Figure 3C). In summary, gene expression across different cell types exhibits distinct enrichments that are functionally relevant to the cells, indicating that various tissues within the seed possess unique expression patterns.

**Figure 3.**
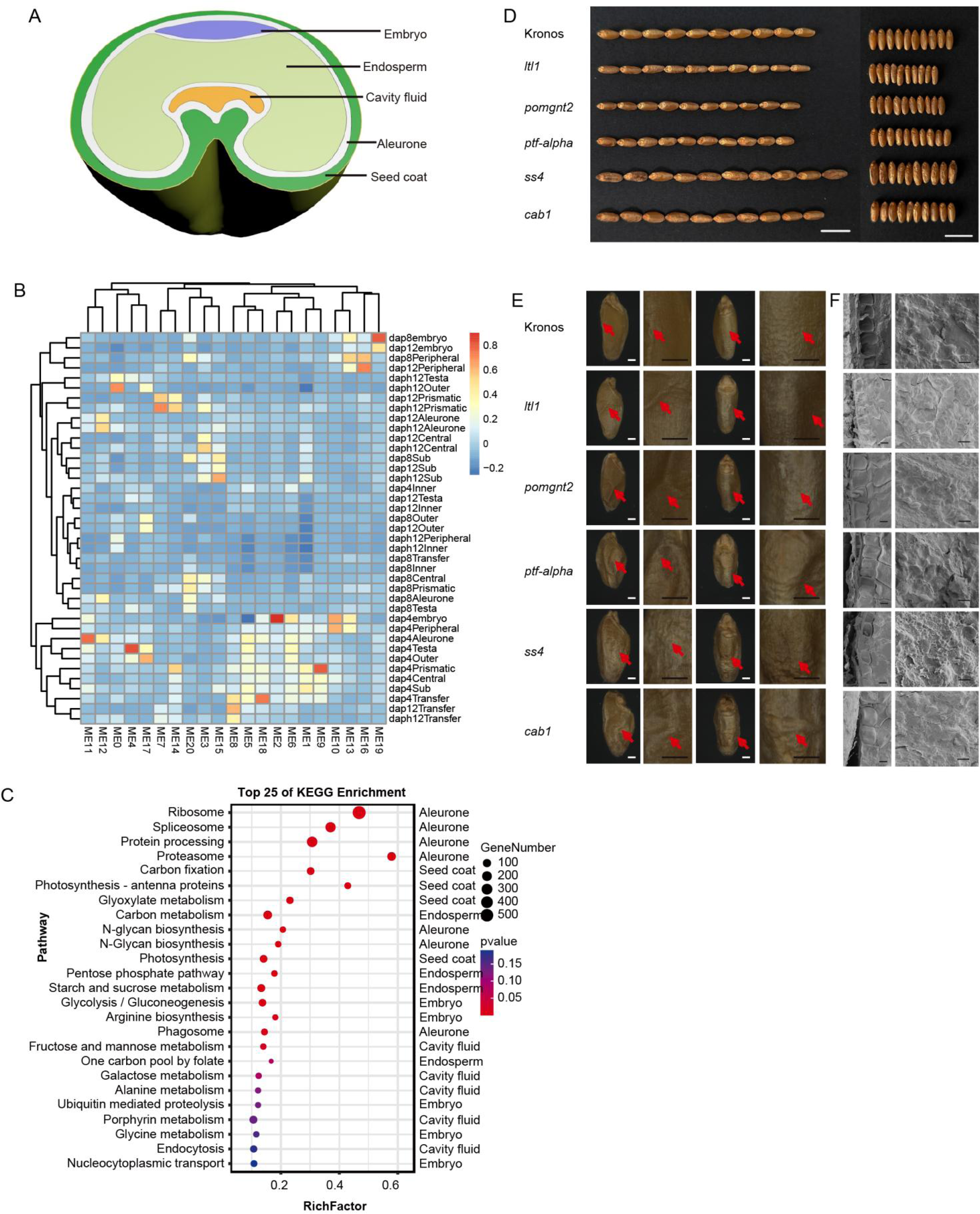
Spatiotemporal co-expression networks for ten grain cell types during development A.The conceptual model for the wheat grain. B. Spatiotemporal co-expression networks for ten grain cell types during grain development. The annotated colours of the columns represent different patterns of co-expression modules and the annotated colours for rows represent development times. C. KEGG enrichment for differentially expressed genes. The enriched KEGG categories were determined using the one-sided version of Fisher’s exact test, followed by the Benjamin-Hochberg correction to obtain adjusted p values for multiple testing. D. Photographs of grain lengths and widths of mutants and wild type (WT). Scale bar = 1 cm. E. Photographs of grain surface. Scale bar =1 mm. Red arrows showed wrinkled seed coat. F. Scanning election microscopic analysis of aleurone layer and endosperm starch. Scale bar =50 μm. Red box showed aleurone layer cell, red arrows showed A-type starch granules, yellow arrows showed B-type starch granules.

We utilized Kronos mutants to validate the functions of representative cell type-specific genes (Figure 3D, Figure S8). The aleurone layer mutant *ltl1* exhibits reduced aleurone cell size and unclear cell edges (Figure 3E). The endosperm cavity mutants *pomgnt2* and *ptf-alpha* display smaller grains with reduced encasement of endosperm starch granules. The endosperm mutant *ss4* shows enlarged grains but with defective endosperm filling, resulting in underfilled grains with an abundance of B-type starch granules. The pericarp mutant *cab1* showed a wrinkled seed coat (Figure 3E). Thus, it is evident that mutations in grain-specific genes affect grain development and filling.

### Transcriptome dynamics at the transition stage during endosperm differentiation

Endosperm development encompasses intricate processes, including cell division, enlargement, and the accumulation of storage compounds such as starch and protein. Understanding the trajectory of wheat endosperm development is essential for unraveling the molecular mechanisms and regulatory networks that control grain filling and seed development. To investigate the shared progenitor cells and the cell fates necessary for their differentiation, we analyzed all major endosperm cell types at various stages. Differentiation was observed at 8 dap and notably increased at 12 dap, especially in the starchy endosperm (Figure 4A-B). The trajectory of cell-type changes was associated with dynamic changes in gene expression, determining cell fates. Thus, we analyzed the top 3000 spatially varying genes correlated with the pseudotime of development and categorized them into four clusters (Figure 4C). We annotated the cell clusters and established a developmental trajectory for the endosperm. Initially, the endosperm cavity, aleurone layer, and peripheral endosperm cells formed a continuous path, with gene expression profiles enriched in ribosomal, DNA replication, and amino acid synthesis processes, indicating active cell division and differentiation. The aleurone layer and external regions of the starchy endosperm (ERS) gradually differentiated towards the distal end of the upper branch, with gene expressions enriched in carbon fixation, and monosaccharide and amino acid metabolism and transport, suggesting their role in nutrient transport. On the lower branch, the endosperm cavity and central endosperm transitioned gradually from the distal end to the central and prismatic endosperm. The gene expression profiles in this path showed enrichment in sucrose and starch synthesis, galactose metabolism, and plant hormone biosynthesis.

**Figure 4.**
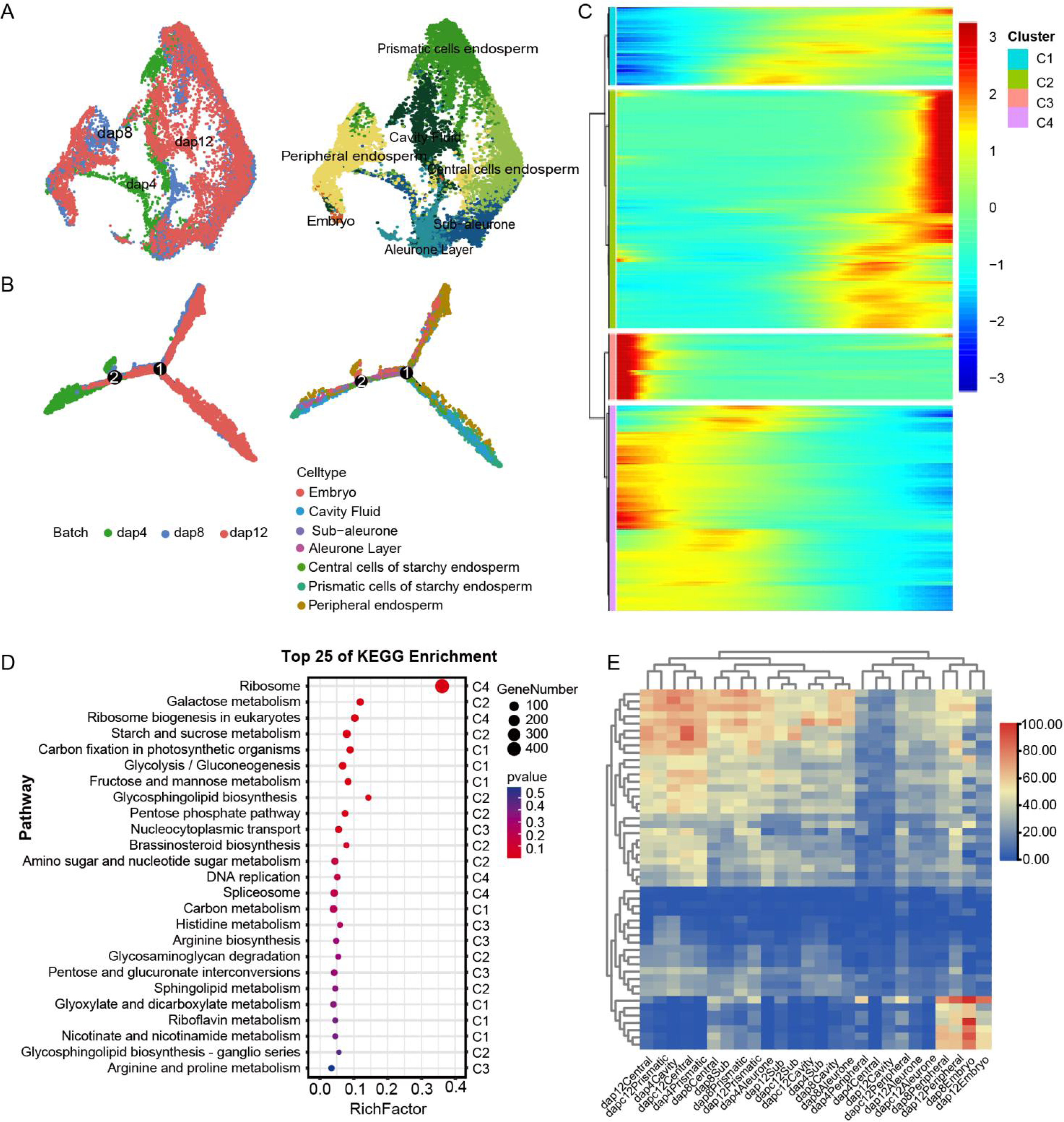
Developmental trajectories of wheat early endosperm. A. Visualization of grain cells via UMAP during the grain development process for merged data (top) and respective data of different radiation times (bottom). B. Visualization of grain development along with pseudotime. C. Expression of the top 3000 gene. D. KEGG enrichment for differentially expressed genes. The enriched KEGG categories were determined using the one-sided version of Fisher’s exact test, followed by the Benjamin-Hochberg correction to obtain adjusted p values for multiple testing. E. Heatmap of the expression of cluster1 and cluster2.

We analyzed gene expression related to carbon fixation, glycolysis, and sugar and starch synthesis in cluster 1 and cluster 2. These genes were highly expressed in the central and prismatic endosperm at 12 dap. The enolase-like isoform (TraesCS5B02G012300) had the highest expression in the prismatic endosperm, indicating its key role in starch synthesis there. Phosphoglycerate kinase (TraesCS6B02G187500) and other sugar synthesis genes were highly expressed in the central starchy endosperm at 8 and 12 dap, suggesting their primary role in nutrient storage in these cells.

The analysis confirmed that endosperm is recognized to undergo dual differentiation pathways, delineated for nutrient transport and storage functions, respectively. These pathways collectively orchestrate grain filling processes, thereby facilitating enhanced nutrient storage within the grain.

### B3 domain-containing transcription factor TaABI3-3B identified through Stereo-seq contribute to grain development

Transcription factors (TFs) play crucial roles in the development of wheat grain by regulating gene expression during various stages. In total, 4,617 TF genes were identified, including 1,535 genes from the A subgenome, 1,520 genes from the B subgenome, and 1,526 genes from the D subgenome. TF genes showed significant differences across cell types different stage (Figure S9). Among the data, 534 transcription factors (TFs) were specifically expressed in the embryo, which is the highest number observed across all cell types. Additionally, we found candidate genes expressed in both the embryo and endosperm, which may influence the differentiation of both tissues. We conducted a subgenomes preference analysis of TFs and found no significant overall gene expression preference (Figure S10). The expressed TF families AP2, ARF, B3, HOME, NAC and WRKY showed embryo-specific enrichment, whereas B3, BZIP, DOF, MYB, NAC and WRKY TFs were enriched in endosperm, and BHLH, BZIP, GRAS and MADS TFs were enriched in the pericarp which is consistent with previous reports (Table S5). We selected six representative gene families for PCA analysis and found that AP2, ARE, HOME, MADS, and WRKY showed little clustering difference in tissues at different stages, whereas B3 displayed distinct clustering at 4 dap, 8 dap, and 12 dap, especially in embryo and endosperm, indicating its divergent biological functions. We conducted a preferential analysis of the B3 gene family and found a B subgenome preference in the endosperm and aleurone layer at 8 dap. However, in the embryo, there was no obvious preference, except for one gene, *TaABI3-3B* (TraesCS3B02G452200), which exhibited higher expression from the B subgenome (Figure S11).

Combine the developmental trajectory data and spatially varying genes in embryo and endosperm, we identified *TaABI3-3B*, acts as a key regulatory gene in embryo and endosperm (Figure 5A). In situ hybridization revealed *TaABI3-3B* specific expressed in early developmental embryos and the surrounding endosperm (Figure 5B). Mutants of *abi3* displayed notable enlargement in embryos and grains size (Figure 5C-D). The phenotypes observed in transgenic lines were consistent with those of mutants. cross-sectional of mutants and transgenic line revealed increased grain area and enlargement of aleurone layer cells (Figure 5F-G), Statistical analysis of yield traits in T3 generation transgenic materials showed increases in grain length, grain width, and hundred-grain weight (Figure 5E), with no adverse effects on normal plant growth, indicating that downregulation of *TaABI3s* expression can enhance wheat yield.

**Figure 5.**
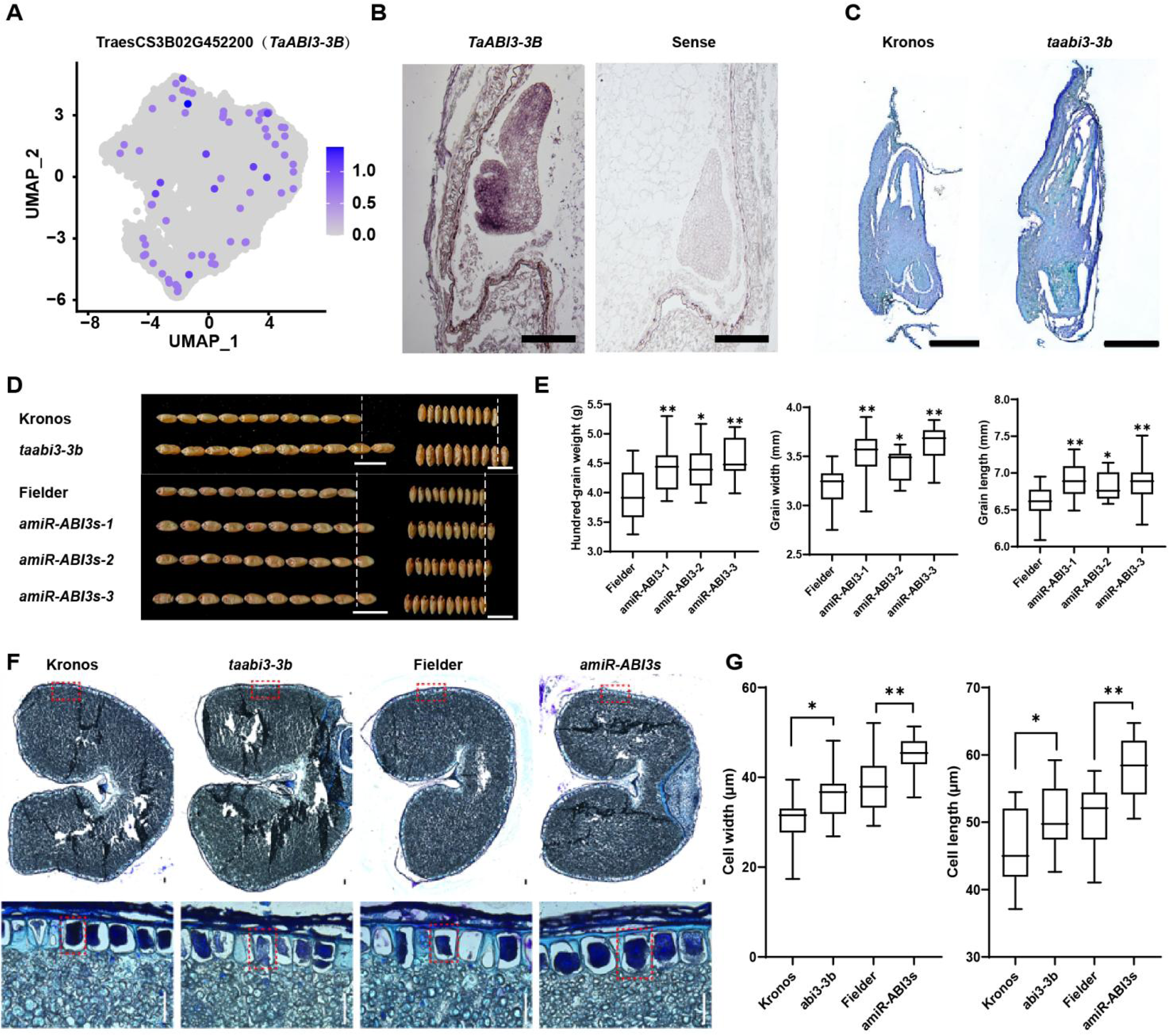
*TaABI3-3B* regulates the grain size of wheat A. Transcript distribution of *TaABI3-3B*. Color bar, normalized UMI counts. B. Spatiotemporal expression pattern of *TaABI3-3B* in dap 10 embryo as indicated by in situ hybridization. Sense probe served as negative control. Scale bars = 100 μm. C. Observation of mature embryo. Scale bars = 1 mm. D. *amiR-ABI3s* and *abi3-3b* mutants increases the grain size in wheat. Scale bars = 1 mm. E. Quantification of grain size related traits between the WT plants and *amiR-ABI3s* lines. Student’s t-test was used to determine the difference significance between *amiR-ABI3s* and WT. *, P ≤ 0.05; **, P ≤ 0.01. F. Representative cross sections of muture grains. Scale bars = 100 μm. The red box shows the location of the enlarged image on the bottom panel. The experiment was independently repeated three times. G. *amiR-ABI3s* and *abi3-3b* mutants significantly increases the AL cell size in wheat. *, P ≤ 0.05; **, P ≤ 0.01.

### Genetic variations in *TaABI3-3B* contribute to with grain weight and quality

To explore the correlation between natural variations in *TaABI3-3B* and grain related traits (grain length, width, weight and quality), the polymorphisms were analyzed in the coding region, 2 kb promoter regions and 2 kb downstream region of *TaABI3-3B* in re-sequencing Watkins collection, which consists of 1,056 hexaploid wheat landraces that represent global wheat diversity (Song et al., 2023). 14 SNPs were found in the promoter of *TaABI3-3B*, as well as 4 SNPs in exon, and 19 SNPs in 3’ downstream region. Based on the SNP, the 1,056 wheat accessions were divided into three groups: 615 with *TaABI3-3B Hap-1* and 150 accessions with *TaABI3-3B Hap-2*, 124 with *TaABI3-3B Hap-3* (Figure 6A). Accessions with *TaABI3-3B-Hap1* displayed higher grain weight, grain length, and grain width and identified as a high-yield haplotype (Figure 6B), while *TaABI3-3B-Hap2* exhibits greater hardness, which facilitates the grain storage. These results indicate that the expression of *TaABI3-3B* contributes to the differences in grain size and quality among wheat varieties. Based on yield data and SNP variation, SNP 4,100 (C>T) of *TaABI3-3B* was identity as key variation site, the C>T change cause the substitution of an acidic amino acid with a neutral amino acid (Glu to Gly), which may affect the function of *TaABI3-3B*.

**Figure 6.**
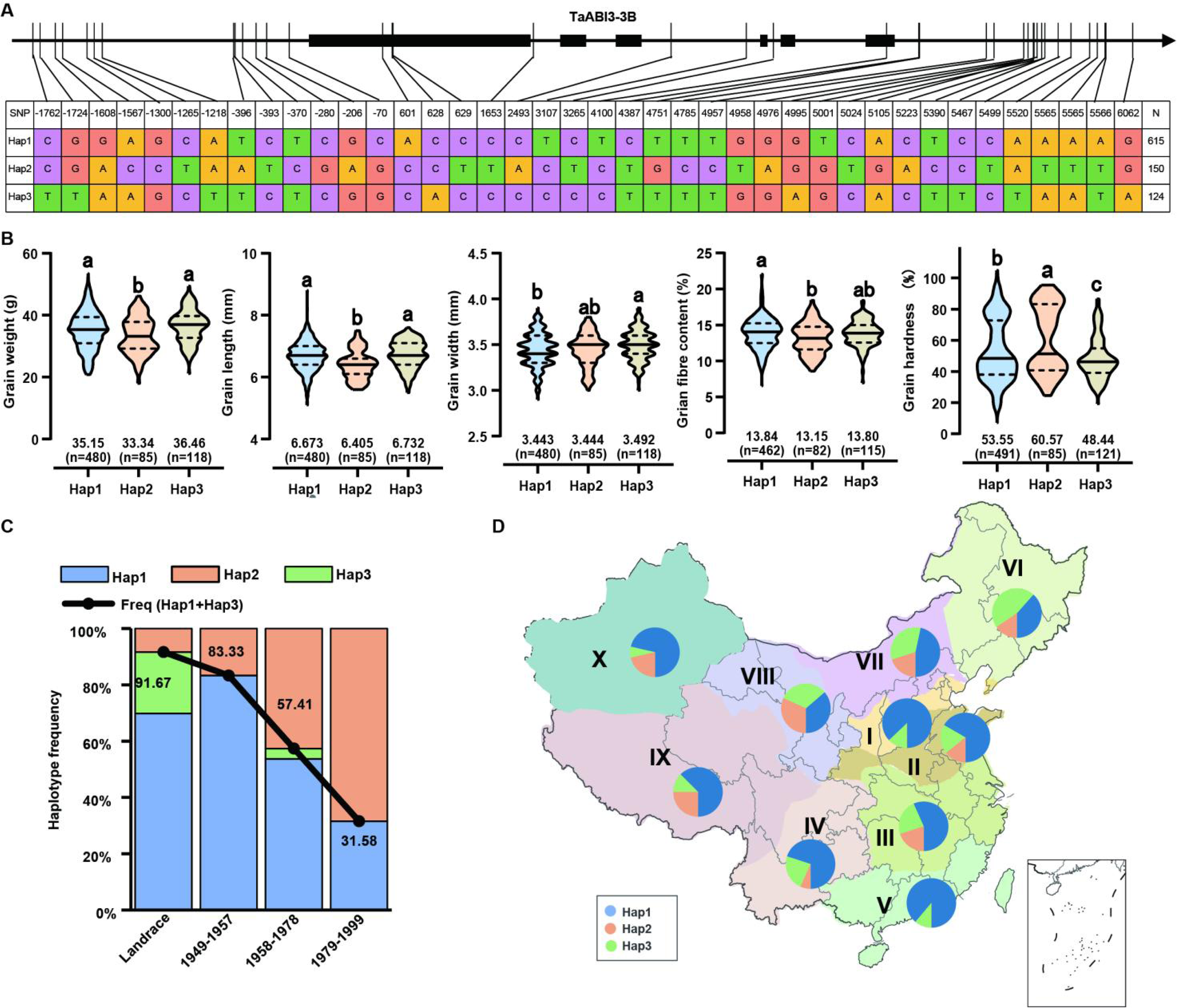
Haplotype analysis of *TaABI3-3B* and breeding selection of elite allele A. Schematic diagram showing the polymorphism for each haplotype of *TaABI3-3B*. The coordinate is related to transcription start site (TSS). B. Violin plot indicating the comparison of grain size related traits among wheat accession with different haplotypes of *TaABI3-3B*. The student’s t-test was used to determine the statistical significance between two groups. C. The percentages of accessions carrying different allele of SNP 4,100 in each categories and during the different breeding process in China. D. The percentage of accessions carrying different allele of SNP 4,100 in each ecological zones of China. The size of pie charts in the geographical map shows the number of accessions, with percentages of the three allele in different colors.

To determine the selection of the high-yield haplotype *TaABI3-3B Hap-1* in China’s wheat breeding process, a detailed analysis was conducted on the SNP 4,100 allele frequency within a mini-core collection (MCC) population of wheat. This investigation revealed that the *TaABI3-3B Hap-2* (C-allele, associated with the haplotype *TaABI3-3B Hap-1* (T-allele), is significantly more prevalent in modern wheat cultivars compared to traditional landraces and cultivars introduced from other countries (Figure 6C). Furthermore, the study highlighted variations in allele distributions across different major agricultural ecological zones in China. Specifically, it was observed that the frequency of the T allele, another variant at the SNP 4,100 site, was higher in Zones I-V compared to other regions (Figure 6D). This geographic distribution indicate that specific ecological conditions may influence the selection and prevalence of different alleles, adapting wheat cultivars to diverse environmental conditions within the country.

## Discussion

Wheat grain development have been intensively studied, including the molecular mechanisms underlying embryo and endosperm development, dynamic regulatory pathways during embryogenesis, as well as nutrient mobilization and storage within the grain. Many techniques such as Laser Capture Microdissection (LCM) and X-ray Micro-Tomography have been employed in the study of grains development (Boudichevskaia et al., 2020; Legland et al., 2023; Xiang et al., 2019). However, there remains a notable gap in research focusing on the comprehensive analysis of whole-seed morphology at the single-cell level in wheat.

In this study, we generated a high-resolution spatial transcriptomics atlas aimed at elucidating gene expression patterns during the early developmental phase comprehensively components. Through this approach we subdivided the grain into 10 discrete cell type. We identified some cell type-specific genes, which are consistent with previously identified endosperm markers from various cereals, such as *TaSS1*, homolog of maize *Shrunken 1 (SH1)* (Chen et al., 2018). *TaRKIN1*, homolog of rice *Sucrose non-fermenting-1-related protein kinas 1b* (*SnRK1b*) (Kanegae et al., 2005). *TaSBE1*, homolog of maize and barley *Starch branching enzyme* (Fu et al., 2023). Meanwhile, we also identified some genes shows different expression pattern and their temporal expression dynamics pattern are differed across species, such as *TaAL9* first detected high expressed at 4 dap at central starchy endosperm and prismatic endosperm, while *HvAL9* in barley expressed exclusively 8 dap at aleurone, which indicate the timing of cell type formation and distinctive gene function differs across cereals. The aleurone layer, sub-aleurone, central cells of starchy endosperm, prismatic cells of starchy endosperm, and ESR markers were already expressed at 4 dap, consistent with previously reported data in barley endosperm, which shows specificity of AL, ETC, and ESR markers were already expressed at 4 and 8 dap (Hertig et al., 2023; Kovacik et al., 2024). This indicates endosperm differentiation is initiated before cellularization.

Previously reported genes involved in grain size regulation and grain quality were found to exhibit cell type-specific expression in our spatial data. For instance, *ATG8a* showed high expression in the pericarp at 4 dap, influencing grain size by regulating the rate of pericarp degradation (Li et al., 2021). PHS was robust expressed at 8 dap and 12 dap in the central endosperm and prismatic cells, which indicates the spatiotemporal specific expression of *PHS1* is crucial for the initiation of B-type granules (Kamble et al., 2023). *MADS29* showed high expression in the cavity fulid at 4 days influence transportation of nutrients into the endosperm and wholly filling of developing grains (Liu et al., 2023). In our spatiotemporal transcriptomics, we have discovered several high-confidence novel candidates with potential regulatory roles. Additionally, natural variations at these gene loci with GWAS analysis (Khan et al., 2022; Yang et al., 2021) and the use of a wheat mutant library to validate gene function would greatly help wheat grain development study and grain yield and quality improvement.

Hexaploid wheat contains three homoeologous chromosome sets, the A, B, and D subgenomes. To investigate if different types of cells within wheat grains has subgenome bias, we analysis various cell types at different time points. We did not observe significant subgenome preference with overall gene expression (Figure S12). Previous study indicates that the biased expression of homoeologs during embryo and endosperm development is associated with differences chromatin accessibility across the A, B, and D subgenomes (Pei et al., 2023). We combined the published genomewide profiling of ATAC-seq data in embryo and endosperm (Zhao et al., 2023) and our spatial transcriptomics data of different tissues to better identify the epigenetic influence during embryo and endosperm. The results showed that chromatin accessibility was associated with the biased expression at A subgenome in the embryo at 8 dap, while there was no clear preference in the endosperm. However, as *TaABI3-3B* is a transcription factor that shows bias towards the B subgenome in both embryo and endosperm, we also analyzed its preference in ATAC and found no significant difference (Figure S12), indicating that *TaABI3-3B* expression is mainly transcriptionally regulated rather than epigenetically modified.

In conclusion, we utilized spatial transcriptomics to identify a large number of genes with spatiotemporal expression specificity. Numerous genes display spatially and temporally precise expression profiles within embryonic tissues. Notably, certain transcription factors exhibit predominant expression within distinct cell types or localized regions of the embryo, indicative of their involvement in modulating cell fate determination and tissue patterning processes (Song et al., 2021). Apply spatial transcriptomics in wheat grain study also has its limitations, particularly when it comes to genes with low expression abundance at the early stage in embryo development. Many gene associated with zygotic activation was mainly observed at dap 2 and dap 4, including cell cycle and cytokine signaling genes, but sharply decrease at dap 8 and dap12 (Xiang et al., 2019; Zhao et al., 2023), detect the low expressed genes remains challenging. Thus, integrating spatial transcriptomics data with complementary techniques such as single-cell RNA sequencing or immunohistochemistry can help overcome some of these limitations and provide a more comprehensive understanding of gene expression patterns within tissues.

## Materials and methods

### Plant growth and tissue preparation

The wheat cultivar JM22 was used in this study. The plants were grown in the greenhouse at 23 °C/18 °C day/night, under long day conditions (16 h light/8 h dark). The grain were harvested at 4, 8 and 12 dap.

### Grain fixation, staining and imaging

Wheat grains at 4, 8, and 12 dap were prepared as follows: soaked in a 75% optimal cutting temperature compound (OCT, Sakura Finetek Europe B.V.) solution under vacuum for 5 minutes, then embedded in OCT and stored at -80 ℃. Once equilibrated to -20 ℃, samples were sliced into 10 μm thick sections. Quality-approved samples were mounted onto Baichuang S1000 chips, treated with methanol fixation, and stained with TBO for imaging.

### Library construction, sequencing and expression atlas analyses

Tissue sectioning, TBO staining, imaging, and initial permeabilization were conducted following the BMKMANU S1000 Tissue Optimization Kit user guide (BMKMANU, ST03003). The secondary permeabilization (8 minutes) and library construction were also performed according to the user guide. The Illumina library was sequenced using the Illumina NovaSeq platform. Raw sequencing data were mapped to the wheat reference genome (IWGSC RefSeq v1.1) using STAR v2.5.3a (Dobin et al., 2012) with default parameters. The image, adjusted by BSTViewer V1.42, along with the corresponding level 4 (27 μm) matrix, was used for downstream analysis. Quality control and normalization of gene expression matrixes for respective samples were performed using Seurat V 4.3.0.1 (Butler et al., 2018) with parameters set to min.cells=5, min.features=100. Batch effect correction was carried out with IntegrateData function in the Seurat package with 3000 anchors. The scaled data were further used for dimension reduction and clustering with 30 principal components and resolution of 0.5. The cell identities defined by spatial information were used for annotation of each cluster. Specific expression genes in respective clusters were identified with FindAllMarkers with logfc.threshold=0.1, only.pos=TRUE, min.pct=0.01. The expression level for each gene for cell types of four samples was identified by two methods: 1, mean=sum of normalized UMI counts/ number of cells, and 2, expression proportion=number of cells with expression/number of cells * 100. The resulting expression matrixes were analyzed using the WGCNA (v.1.69) pipeline with the default filtration process for dentification spatially co-expressed gene modules (Langfelder and Horvath, 2008).

### Construction of embryo and endosperm cell trajectories

The Seurat objects of embryo and endosperm cells were extracted for reconstruction of cell atlases with the same pipeline described above. Two thousand of highly variable genes were identified. The expression matrix was further analyzed and ordered by the identified highly variable genes using monocle2 v2.14.0 (Qiu et al., 2017). Dimension reduction was performed with DDRTree methods. Differential expression genes along the pseudotime were identified and the top 2k genes were used for another round of dimension reduction.

### Marker genes identification and RNA in situ hybridization

After analyzing the gene expression of cell populations, the top 20 genes in each cell population were specifically expressed in tissues and gene functions, and some genes were used as the maker genes of the cell population. After electron in situ hybridization, these genes were randomly selected to express tissue-specific and functionally related genes for in situ hybridization validation. Using gene specific fragments as templates, synthesize DIG labeled probes. Fix the developing grain with formalin-aceto-alcohol solution (50%), dehydrated in a series of ethanol concentrations and cleared in histoclear, embedded in paraplast (Sigma, P3558) and sectioned to a thickness of 7 μm. The following step of RNA hybridization, immunologic detection and signal capture with the hybridized probes were compiled as described previously (Meng et al .,2017)

### Scanning electron microscopy

The grains were broken transversely, mounted on aluminum stubs, coated with gold. Hitachi S-3400N (Hitachi, Tokyo, Japan) scanning electron microscope was used to observe the samples.

### KEGG Enrichment Analysis

The enrichment level of differentially expressed genes (DEGs) in the KEGG pathways (http://www.genome.jp/kegg/) was determined using the KOBAS (3.0) software. with a threshold of FDR ≤ 0.05 considered for defining significantly enriched pathways.

### Creation of transgenic plants of *TaABI3s*

To knock down *TaABI3s* in wheat cv. Fielder, the oligos for amiRNA 5’-AAAATCGGTACCGCATGCTT-3’ located in the third exon was used. The amiRNA vector was conducted following the methods as previously reported (Park et al., 2009). The amiR vectors were introduced into hexaploid wheat (cv. Fielder) via *Agrobacterium*-mediated transformation as described by Ishida et al. (2015). PCR, herbicide (glufosinate) spraying and a QuickStix Kit for bialaphos resistance (bar) were used to verify the positive transgenic plants.

## Figure Legends

Supplementary Figure 1. Overview of Spatial Transcriptomics experiment on wheat develoment grains.

Supplementary Figure 2. The analysis pipeline for spatial transcriptomic study.

Supplementary Figure 3. The density of expressed genes and transcripts in a spot.

A. The number of expressed genes in a spot. The y-axis represents the number of genes expressed in a spot. The y-axis represents the number of genes expressed in a spot. B. The spatial distribution of expressed genes on the sample sections. The color changes gradually from green, yellow to red when the density gradually increases. C. Pearson correlation among UMI and genes.

Supplementary Figure 4. Spatial visualization of the unbiased spot clustering for 4, 8 and 12 dap wheat sections. Merged bright field image and spatial clusters of other three sections.

Supplementary Figure 5. The defined clusters after dimensional reduction.

A. The 6 clusters from dimensional reduction are mapped back to tissue sections, showing their location on the aleurone layer. The clusters are distinguished by different colors. B. The 5 clusters from dimensional reduction are mapped back to tissue sections, showing their location on the sub aleurone. The clusters are distinguished by different colors. C. The 9 clusters from dimensional reduction are mapped back to tissue sections, showing their location on the cavity fluid. The clusters are distinguished by different colors.

Supplementary Figure 6. Representatives of electronical RNA in situ hybridization using known markers that are consistent with the previous studies (Xiang et al., 2019). We selected two markers from three tissue as following due to unavailability from other research: *TaBBM* (TraesCS6B02G091800) in embryo; *TaBGLU16* (TraesCS2A02G038400) and *TapsaD* (TraesCS5D02G468400) in seed coat; *TaAMY2* (TraesCS2B02G004100), *TaCYP71D7* (TraesCS4A02G363000) and TraesCS4A02G363000 in endosperm.

Supplementary Figure 7. Representatives of electronical RNA in situ hybridization using newly defined markers.

We selected one marker from each compartment as following: lipid transfer protein TraesCS3A02G344700 in pericarp, non-specific lipid-transfer protein TraesCS5B02G531100 in aleurone layer, protection and storage gene TraesCS7A02G183600 in sub-aleurone layer, SWEET gene TraesCS7B02G160000 in cavity fluid, transcription factors TraesCS6D02G258400 in embryo, gibberellin-regulated protein 1 TraesCS5D02G408100 in ESR, TraesCS5B02G353200 has been selected as the marker gene for this characteristic, alpha-amylase TraesCS4B02G328100 in central cells of starchy endosperm, seed storage-related proteins TraesCS1A02G063100 in prismatic cells of starchy endosperm.

Supplementary Figure 8. Representatives of electronical RNA in situ hybridization using mutants gene.

The aleurone layer mutant *ltl1* (TraesCS1A02G145300), endosperm cavity mutants *pomgnt2* (TraesCS6B02G339100) and *ptf-alpha* (TraesCS6A02G279100), the endosperm mutant *ss4* (TraesCS1A02G353300), the pericarp mutant *cab1* (TraesCS5B02G353200).

Supplementary Figure 9. Heat map showing the expression of TFs in 10 grain cell types across three developmental stages.

Supplementary Figure 10. Balanced homoeologs in TFs expression show unbalance expression patterns in 10 grain cell types across three developmental stages defined by stRNA-seq data. The ternary plot shows expression pattern of detected genes in bulk RNA-seq.

Supplementary Figure 11. Diverse expression patterns of TFs across cell types and stage.

A. TF family PCA in all stage of 10 cell types. B. Balanced homoeologs in B3 family show various unbalance expression patterns in different cell clusters defined by stRNA-seq data. The ternary plot shows expression pattern of detected genes in bulk RNA-seq.

Supplementary Figure 12. Balanced homoeologs in genes expression show unbalance expression patterns in 10 seed cell types across three developmental stages defined by stRNA-seq data. The ternary plot shows expression pattern of detected genes in bulk RNA-seq.

Supplementary Table S1. The statistics of sequencing data.

Supplementary Table S2. The summary of marker genes for 10 cell types from the literature RNA-seq from laser microdissection.

as well as experimental in situ hybridization.

Supplementary Table S3. Molecular marker genes.

Supplementary Table S4. WGCNA module genes.

Supplementary Table S5. All TFgenes and TF family.

Supplementary Table S6. Primers used in this study.

## Author contributions

Y.C. conceived and supervised the study. X.L. performed tissue sectioning, spatial transcriptomes sequencing experiments and analyzed the data; X.H. conducted bioinformatic analysis and presented the data. Y.W generated transgenic lines, investigated phenotypes, and performed molecular experiments; X.L. and Y.W. did in situ hybridization; D.W. did the haploid and selection analysis; J.W. provided Kronos mutants; X.L.and Y.C. wrote the manuscript; J.W., X.L. and K.C. revised the manuscript; All authors discussed the results and commented on the manuscript.

## Acknowledgements

We thank Dr. Daolin Fu of Shandong Agriculture University for providing Kronos mutants which were originally produced by Dr. Jorge Dubcovsky’s lab at the UC Davis. This work was supported by the Key R&D Program of Shandong Province, China (ZR202211070163), the Taishan Scholars Program (tsqn202103162) and the Natural Science Foundation of Shandong Province (ZR202102190348) to Y.C. and (ZR2021MC041) to X.L.

